# Variation of mutational burden in healthy human tissues suggests non-random strand segregation and allows measuring somatic mutation rates

**DOI:** 10.1101/332734

**Authors:** Benjamin Werner, Andrea Sottoriva

## Abstract

The immortal strand hypothesis poses that stem cells could produce differentiated progeny while conserving the original template strand, thus avoiding accumulating somatic mutations. However, quantitating the extent of non-random DNA strand segregation in human stem cells remains difficult *in vivo*. Here we show that the change of the mean and variance of the mutational burden with age in healthy human tissues allows estimating strand segregation probabilities and somatic mutation rates. We analysed deep sequencing data from healthy human colon, small intestine, liver, skin and brain. We found highly effective non-random DNA strand segregation in all adult tissues (mean strand segregation probability: 0.98, standard error bounds (0.97,0.99)). In contrast, non-random strand segregation efficiency is reduced to 0.87 (0.78,0.88) in neural tissue during early development, suggesting stem cell pool expansions due to symmetric self-renewal. Healthy somatic mutation rates differed across tissue types, ranging from 3.5×10^−9^ /bp/division in small intestine to 1.6×10^−7^/bp/division in skin.

**Author Summary:** Cairn proposed in 1975 that upon proliferation, cells might not segregate DNA strands randomly into daughter cells, but preferentially keep the ancestral (blue print) template strand in stem cells. This mechanism would allow to drastically reduce the rate of mutation accumulation in human tissues. Testing the hypothesis in human stem cells within their natural tissue environment remains challenging. Here we show that the patterns of mutation accumulation in human tissues with age support highly effective non-random DNA strand segregation after adolescence. In contrast, during early development in infants, DNA strand segregation is less effective, likely because stem cell populations are continuing to grow.

## Introduction

The immortal DNA strand hypothesis, originally proposed by Cairns in 1975, poses that adult mammalian stem cells do not segregate DNA strands randomly after proliferation [1]. Instead, stem cells might preferentially retain the parental ancestral strand, whereas the duplicated strand is passed onto differentiated cells with limited life span (Figure 1). In principle, such hierarchical tissues could produce differentiated progeny indefinitely without accumulating any proliferation-induced mutations in the stem cell compartment [2,3]. Experimental evidence supporting this hypothesis comes from BrdU stain tracing experiments both *in vitro* and *in vivo* [4-7]. Evidence from spindle orientation bias in mouse models of normal and precancerous intestinal tissue corroborated these findings, suggesting that strand segregation is then lost during tumourigenesis [8]. However, many of the experiments suffer from uncertainties in stem cell identity and a definite mechanism of strand recognition remains unknown [9]. Hence why Cairns hypothesis remains controversial [10].

**Figure 1:**
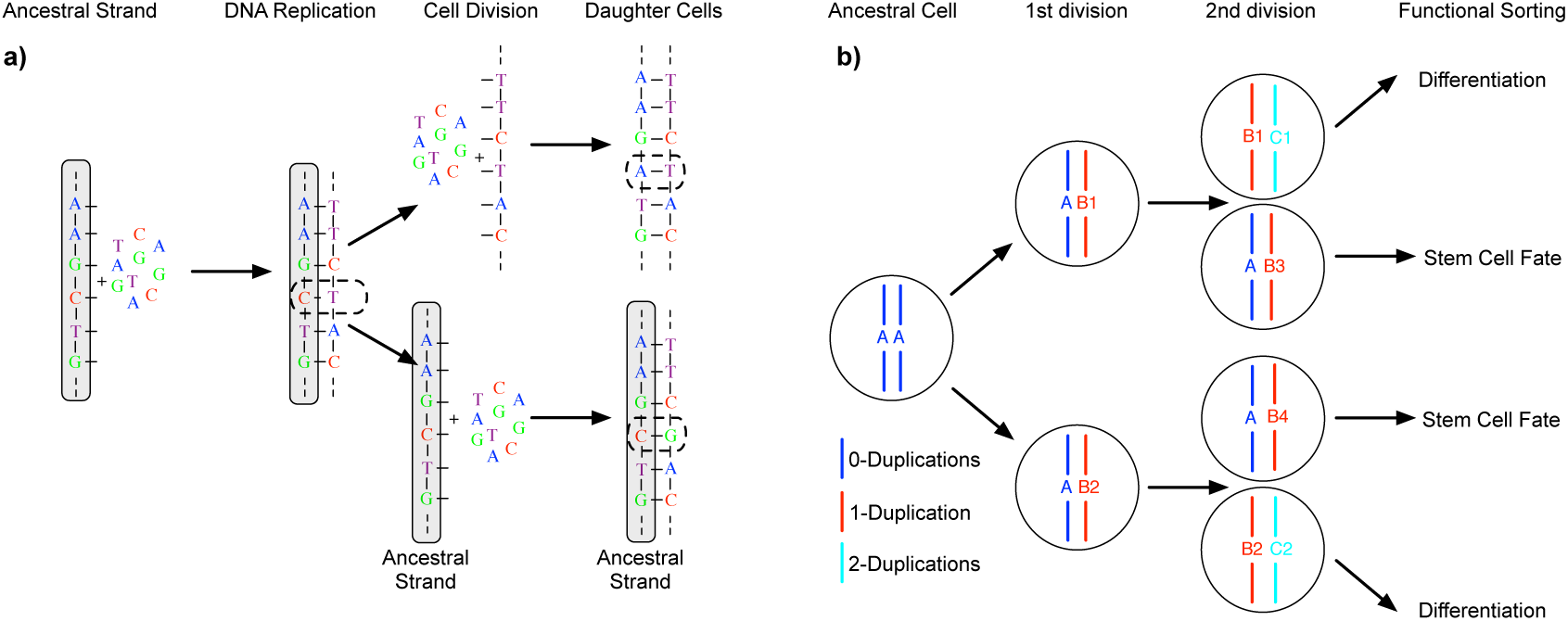
The Immortal DNA strand hypothesis. **a)** During replication of the ancestral DNA strand, errors (dashed line) might occur. If these errors are not corrected by intrinsic DNA repair mechanisms, they become permanently fixed in daughter cells after the next cell division. However, the original ancestral strand is still present and can provide the blue print for additional non-mutated copies of DNA. **b)** In principle, a stem cell driven tissue allows for non-random DNA strand segregation. Preferentially segregating ancestral DNA strands into stem cells and duplicated strands into differentiated cells with limited life span can drastically reduce the accumulation of somatic mutations in tissues.

Orthogonal studies based on the expected accumulation of somatic mutations in healthy human tissues have argued against the immortal strand hypothesis [11,12]. However, the mere accumulation of somatic mutations in healthy tissue neither supports nor negates the immortal strand hypothesis *in vivo*. Here, we show that measuring the change of the mutational burden and, most crucially, the change of the *variance* of the mutational burden with age allows determining the probability of DNA strand segregation and the per cell mutation rate in healthy human tissues. First, we outline the approach and then apply it to genomic data from healthy human colon, small intestine, liver, skin and brain tissue. The data comes from four recent independent studies on mutational burden in healthy tissues [13-16], which contain information on in total 39 individuals at different ages and analysed genomes of 341 single cells. We find evidence for non-random strand segregation in all adult tissues and significant differences in somatic mutation rates between tissues, but less prominent strand-segregation in brain tissue during early development.

## Results

### The expected change of mean and variance of mutational burden with age

We describe the accumulation of mutations with time in hierarchically organised human tissues by a stochastic mathematical and computational model, Figure 1. A detailed description and derivation of all equations is provided below (Materials and Methods). Briefly, our model considers a constant number of *N* stem cells that contribute to tissue homeostasis. Stem cells divide with a certain constant rate *λ*, e.g. once every week or month. During each division, the parental DNA strand is copied and *χ* novel mutations might occur on the daughter strand. Here *χ* is a random number that follows a Poisson distribution with mutation rate *µ* per bp/division and genome size *L*. Cell fate is also probabilistic in our model. Cells with the parental strand will keep a stem-cell fate with probability *p*, e.g. for *p* = 1 they will always remain stem cell, or differentiate otherwise, e.g. for *p* = 1/*2* cell fate decisions are purely random (coin flip). We can understand the probability *p* as the probability of non-random strand segregation, e.g. *p* ≈ 1 suggest highly non-random strand segregation, whereas *p* = 1/*2* corresponds to random strand segregation.

With this model, we can describe the accumulation of mutations over time explicitly (see Materials and Methods for more details). Assuming the mutation rate *µ* as well as the cell proliferation rate *λ* to be constant, we find that both the mutational burden 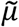 as well as the variance of the mutational burden *σ*^2^ are predicted to increase linearly with time *t*:

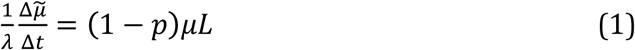

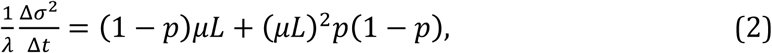

see Materials and Methods for a detailed derivation and Figure 2 for a verification by individual based computer simulations. However, the rates by which the mutational burden and the variance of the mutational burden increase over time depend differently on the mutation rate *µ* and the non-random strand segregation probability *p*. This allows us to independently measure the mutation rate *µ* and the non-random strand segregation probability *p* via:

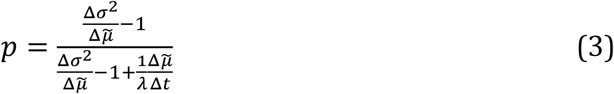

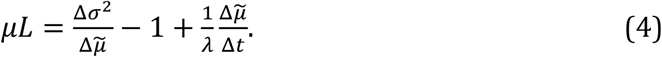

**Figure 2:**
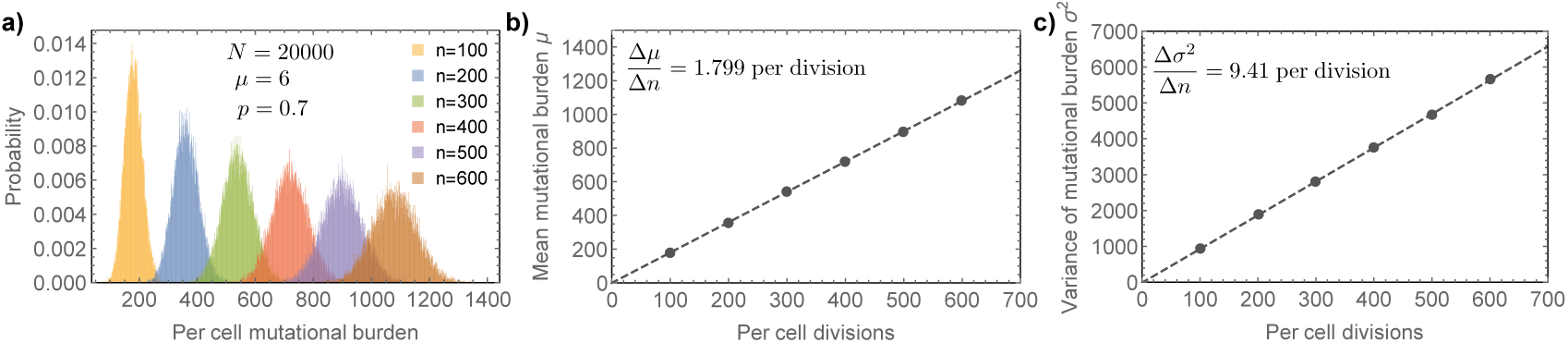
Predicted mutational burden in individual stem cells with age. **a)** We show simulated stochastic mutation accumulation in a stem cell population of constant size. Here *N* = 20,000 stem cells segregating DNA strands with probability *p* = 0.7 and a mutation rate of *µ* = 6 per cell division (corresponding to a mutation rate of *µ* = 10^−9^ per bp per cell division). **b)** Mutational burden and **c)** variance of the mutational burden increase linear. Linear regression (dashed lines) gives 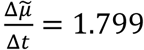 and 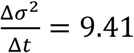. The expected exact values based on above parameters and equation (1) and (2) are 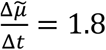 and 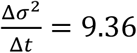. Equation (3) and (4) yield for the strand segregation probability *p* = 0.702 and for the mutation rate *µ* = 6.03, (exact values imposed on the simulation were *p* = 0.7 and *µ* = 6).

Importantly, measuring the change in mutational burden 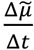 and variance 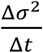 over time in combination with equations (3) and (4) determines the mutation rate *µ* (per cell division) and the non-random strand segregation probability *p* for healthy tissues.

### Measured mean and variance of mutational burden with age from sequencing data

In a recent publication Blokzijl and colleagues [13] measured mutation accumulation in healthy colon, small intestine and liver tissue by whole genome sequencing multiple single stem cell derived organoids of healthy donors of different ages. In addition, Martincorena and colleagues [14] measured mutational burden in multiple skin samples of four individuals with ages between 58 and 73 years. Furthermore, two recent publications [15,16] performed large-scale single cell whole genome sequencing of neurons at different ages. In the experiments by Blokzijl and colleagues [1,13], they isolated single cells and expanded those into organoids. These cells can therefore be thought of as tissue specific stem cells. In contrast, the other experiments [2,3,14-16] do not directly measure mutational burden in stem but more differentiated progenitor cells. However, compared to the total number of cell divisions in the tissue, the number of divisions separating stem and progenitor cells is neglectable.

These datasets enable measurements for the change in mutational burden 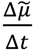 and the variance 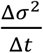 of the mutational burden with age in those healthy human tissues, see Figure 3 & 4. Equations (3) and (4) have a single undetermined parameter, the stem cell proliferation rate *λ*. Strictly speaking, they therefore only provide possible ranges for the mutation rate and the strand segregation probability. However, the possible ranges are narrow for any biologically meaningful stem cell proliferation rate, see Figure 5.

**Figure 3:**
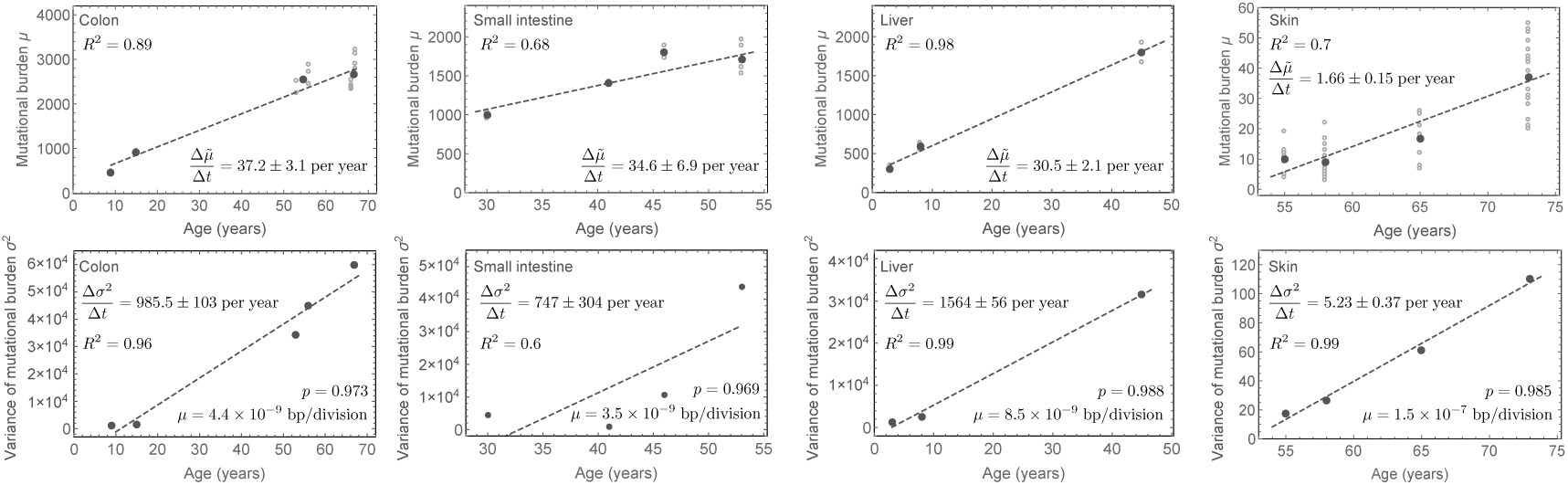
Mutational burden and variance in healthy human tissues. Mutational burden and variance of the mutational burden in colon, small intestine liver and skin tissue in healthy adult humans of different ages, data taken from [13,14]. Open circle represent mutational burden of single cells, whereas dark grey dots represent the mean mutational burden or variance respectively. In all cases, the data well supports our expectation of a linearly increasing mean and variance with age. Linear regressions (dashed lines) give estimates for the change of the mutational burden and the variance with age, see main text (uncertainties represent standard errors). Equations (3) and (4) then allow to estimate the non-random strand segregation probability as well as the per-cell mutation rate per cell division. In all cases, the probability of non-random strand segregation is high (median: *p* = 0.979 (0.97,0.99)), whereas the mutation rate per cell division varies between tissues and is highest in skin, see insets and main text for tissue specific estimates.

**Figure 4:**
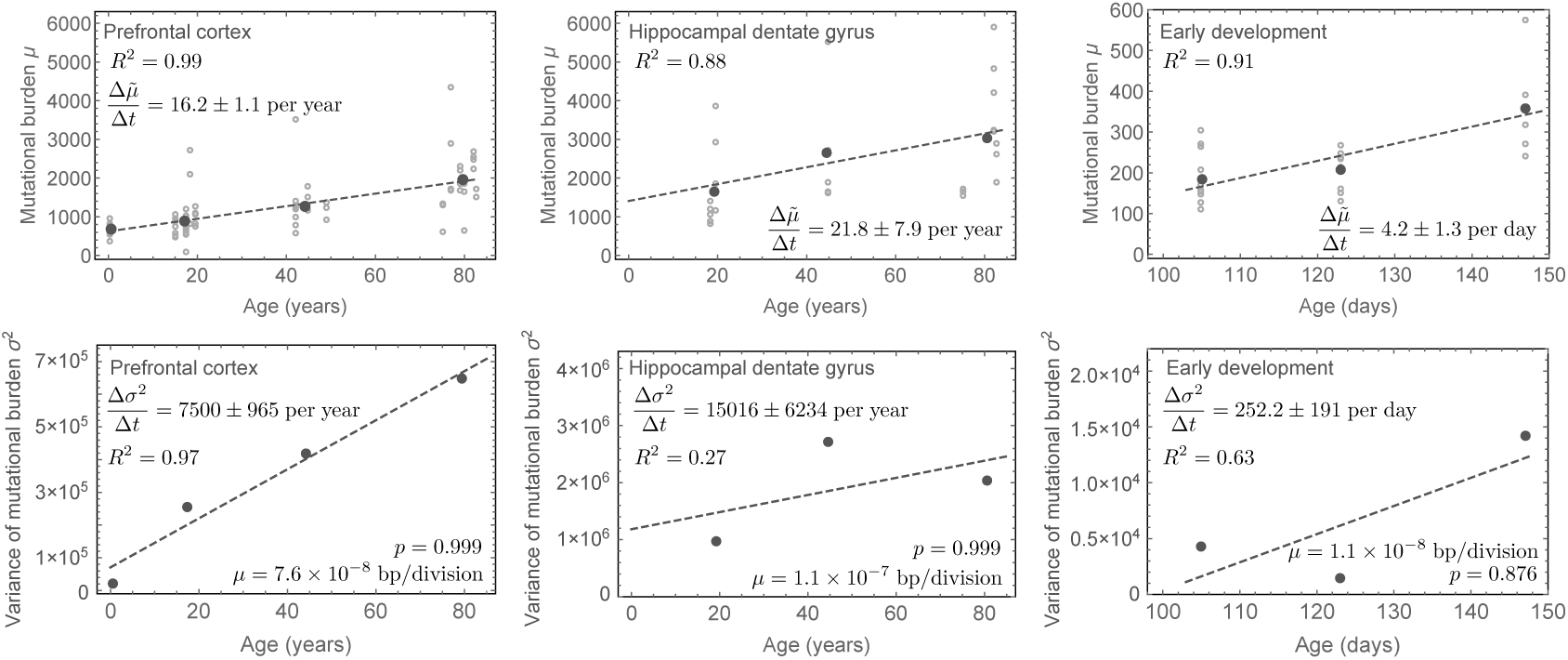
Mutational burden and variance in human neurons during early development and adulthood. Mutational and variance measured from single whole genome sequencing of neurons in the prefrontal cortex and the hippocampus dental gyrus [15] as well as in single neurons during early childhood development after birth [16] (uncertainties represent standard errors). Mutation accumulation in early childhood is highly increased compared to adulthood. However, the per-cell mutation rate per division appears higher in adulthood. The non-random strand segregation in contrast is with *p* = 0.999 (0.998; 0.9993) extremely high in adults, whereas with *p* = 0.876 (0.78; 0.88) it is lower in early childhood. This can be understood as a consequence of cell population expansions due to symmetric self-renewals in early childhood. For details of mutation rate estimates, see the main text.

**Figure 5:**
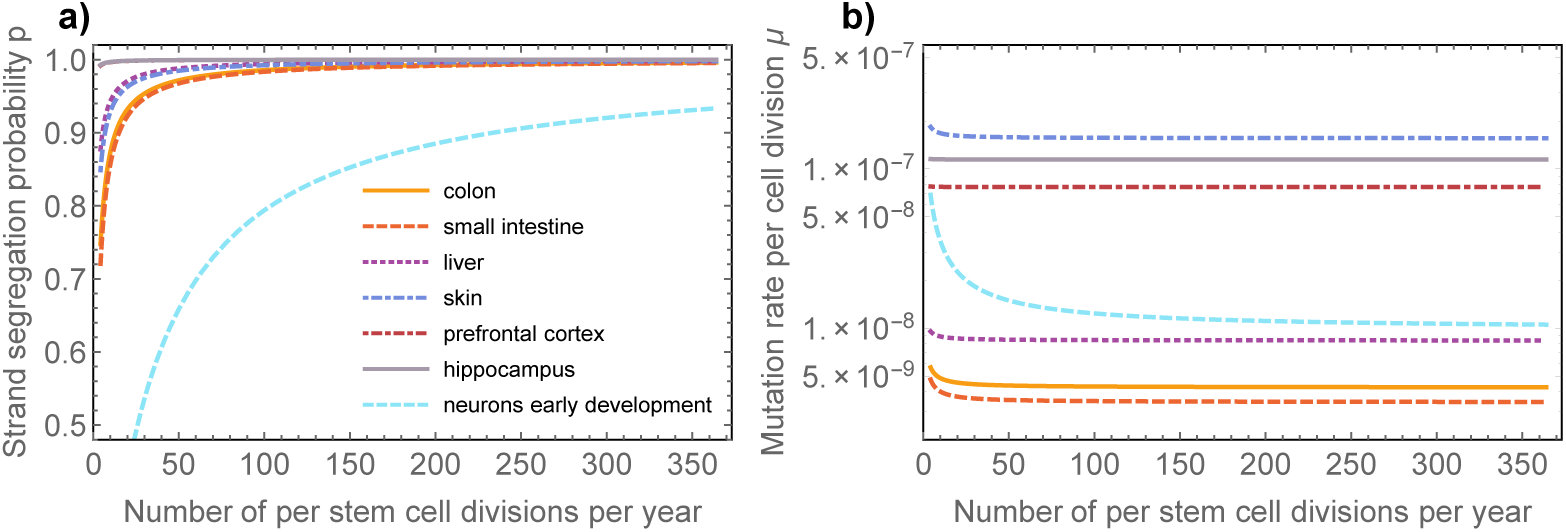
Dependence of parameter inferences on stem cell proliferation rate. Inferences of **a)** the DNA strand segregation probability and **b)** mutation rate per cell division are robust against wide ranges of the stem cell proliferation rate *λ*. If stem cells divide once per week this implies (Equation (3)) for the probability of DNA strand segregation in colon: *p* = 0.973 (0.892; 0.996), small intestine: *p* = 0.969 (0.877; 0.995), liver: *p* = 0.988 (0.952; 0.998), prefrontal cortex: *p* = 0.999 (0.997; 0.9999), hippocampal dentale gyrus: *p* = 0.999 (0.997; 0.9999), skin: *p* = 0.985 (0.94; 0.998). Numbers in brackets correspond to the range of the DNA strand segregation probabilities for stem cell replication rates between once per month and every day respectively. In contrast for neurons during early development we find: *p* = 0.876 (0.67; 0.96) if cells divide every 48h (number in brackets correspond to cell divisions once per week and twice a day respectively). Based on Equation (4) we find for the *in vivo* mutation rate per base pair per cell division in colon: *µ* = 4.37 (4.27; 4.77)× 10^−9^, small intestine: *µ* = 3.54 (3.45; 3.91)×10^−9^, liver: *µ* = 8.48 (8.39; 8.8)×10^−9^, prefrontal cortex: *µ* = 7.68 (7.67; 7.7)×10^−8^, hippocampal dentale gyrus: *µ* = 1.14 (1.14; 1.15)×10^−7^, neurons during early development: *µ* = 1.23 (1.02; 1.47)×10^−8^ and skin: *µ* = 1.57 (1.56; 1.65)×10^−7^.

### Estimations of non-random strand segregation probability in healthy human tissues

For all tissues, the experimental observations confirm our expectation of a linearly increasing mean and variance of the mutational burden. Using linear regression on the data in [4-7,13-16], we find for colon that the change in mutational burden over time was: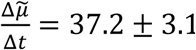, for small intestine: 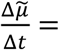 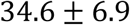, for liver: 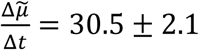, for prefrontal cortex: 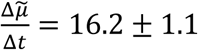 and for hippocampal dentate gyrus: 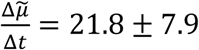 mutations per whole genome per year. We found for skin: 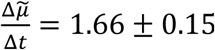 mutations per 0.69 Mb per year. We found for neurons during early development: 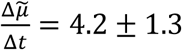 mutations per whole genome per day. Uncertainties here are standard errors. Similarly, for the change of variance we found for colon: 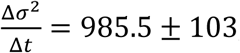, for small intestine: 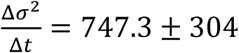, for liver: 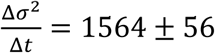, for prefrontal cortex: 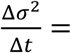 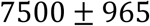, for hippocampal dentate gyrus: 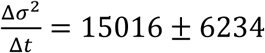 mutations per whole genome per year, for skin: 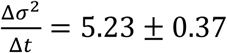 mutations per 0.69 Mb per year and for neurons during early development: 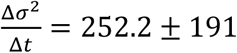 mutations per whole genome per day (Figure 3 & 4).

If stem cells divide once per week this implies (Equation (3)) for the probability of DNA strand segregation in colon: *p* = 0.973 (0.971; 0.974), small intestine: *p* = 0.969 (0.966; 0.97), liver: *p* = 0.988 (0.987; 0.989), prefrontal cortex: *p* = 0.999 (0.998; 0.9993), hippocampal dentale gyrus: *p* = 0.999 (0.998; 0.9998), skin: *p* = 0.985 (0.983; 0.987). In contrast for neurons during early development we find: *p* = 0.876 (0.78; 0.88) if cells divide every 48h. Numbers in brackets correspond to the range of the DNA strand segregation probabilities given the upper and lower bound of the error estimates of the linear regressions. Dependencies of the estimates on the proliferation rate can be found in the caption of Figure 5.

This suggests highly effective non-random DNA strand segregation in human adult stem cells and is in line with previous observations of predominantly asymmetric stem cell divisions [6-8,17,18]. It would require extreme stem cell proliferation rates of approximately one division per stem cell per year for the data to be consistent with solely random strand segregation (*p* = 0.5), Figure 5. This is an unlikely scenario as all tissues analysed here are thought to have high stem cell proliferation rates [9,19,20]. Interestingly, during development non-random DNA strand segregation is less prominent. One explanation is an expanding stem cell population due to symmetric stem cell self-renewals during early development[1,10], which also would explain the increased accumulation of mutations early, as well as typical increased telomere shortening early in life [11,12,21].

#### Measurements of somatic mutation rates in healthy human tissues

Based on Equation (4) we find for the *in vivo* mutation rate per base pair per cell division in colon:*µ* = 4.37 (4.26; 4.46)×10^−9^, small intestine: *µ* = 3.54 (*2*.61; 4.17)×10^−9^, liver: *µ* = 8.48 (8.22; 8.77)×10^−9^, prefrontal cortex: *µ* = 7.68 (7.18; 8.12)×10^−8^, hippocampal dentale gyrus: *µ* = 1.14 (1.04; 1.68)×10^−7^, neurons during early development: *µ* = 1.23 (0.43; 1.52)×10^−8^ and skin: *µ* = 1.57 (1.54; 1.63)×10^−7^. The ranges of these values agree with a recent estimate of the somatic mutation rate in human fibroblasts [13-16,22] and are one to two orders of magnitude larger than germline mutation rates [13,23,24]. However, our method does not require precise estimates of the total number of cell divisions since conception (Figure 5). We find surprising differences in the somatic mutation rates across tissue types that cannot be explained by for example different stem cell proliferation rates alone. The mutation rate estimate in skin is particularly high. This might be due to the nature of the samples used by Martincorena and colleagues [14], as the mutational burden was measured in eye lids of individuals that were exposed to high levels of UV radiation for decades. It is plausible that this contributed to the very high mutation rate estimate. It remains to be seen, if these differences across tissues prevail for denser sampling in more individuals.

#### Explaining strand segregation in terms of symmetric stem cell divisions

Our analysis suggests in general highly effective non-random DNA strand segregation in human colon, small intestine, liver, skin and brain. However, approximately 1% to 5% of divisions in adults do not seem to segregate strands properly and stem cells accumulate additional mutations over time. The reason for this improper segregation could be either wrongly segregated strands during an asymmetric stem cell division or the loss of a stem cell by either a symmetric stem cell differentiation or cell death followed by a symmetric stem cell self-renewal. Arguments are made for both symmetric and asymmetric stem cell divisions in human tissues [15,16,25-28]. We wondered if our approach provides a mean to distinguish both possibilities. We therefore implemented stochastic simulations of mutation accumulation in either asymmetric dividing stem cell populations with imperfect strand segregation or a stem cell population with a mix of symmetric and asymmetric divisions (SI Figure 1). Both scenarios lead to linearly increasing mean and variance of the mutational burden, with small differences in the actual rates. However, as predicted, the ratio of the variance and the mean 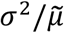 are in both scenarios independent of time and on average the same (see also Equation (S10)). Interestingly, the distribution of 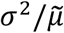 differs. Whereas the variance of the distribution of 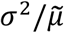 increases with time for symmetric stem cell divisions, it approximately remains constant for asymmetric stem cell divisions. However, measuring this effect reliably would require measuring the mean and variance of the mutational burden in many more independent samples of many more healthy humans of different ages than the currently available datasets. Hence, lack of resolution in currently available data precludes us to determine the cause of imperfect strand segregations. However, this effect might provide a future mean to quantitate the amount of symmetric self-renewal in human stem cell populations.

## Discussion

Stem cells in fast proliferating healthy adult tissues such as colon have been reported to accumulate approximately 40 new mutations per year [13] (Figure 3). However, if mutation rates are in the order of 10^−9^ per base pair per cell division, which seems to be the current consensus and agrees with our measurements here, and the human genome consists of 6×10^9^ base pairs, this would on average only allow for 6 to 7 divisions per stem cell per year. This is in contradiction to current measures on stem cell turnover rates in for example healthy colonic crypts [19,29]. This discrepancy is resolved by non-random strand segregation, where many stem cell proliferations would not induce novel mutations on the stem cell level and the effective observed mutation accumulation on a population level can remain low despite high stem cell turnover rates. A clear molecular mechanism of strand recognition remains unknown. However, direct and indirect evidence to which our observations may contribute increasingly hint on the importance of strand segregation to maintain genomic integrity within healthy human tissues.

Our joined inference of mutation rate and strand segregation probability also reveals that mutation rates per cell division are likely higher than was assumed in previous studies [11]. We therefore find stronger signals of strand segregation in human sequencing data than was thought previously [11]. Our inference neglects the effects of cell-division independent mutations that may contribute to mutational burden in tissues at a low rate. This can lead to an underestimation of the true strand-segregation probability as well as the per-cell mutation rate in human tissues, see SI Figure 2.

A loss of strand segregation in stem cells implies a 50 to 100 times increased effective mutation rate on the cell population level without any other changes to the intrinsic DNA repair machinery. In a non-homeostatic setting, such as a growing tumour, in which the number of self-renewing cells (whether they are all or only a subset of cells) increases, the rate of random strand segregation events is much higher. This effect may contribute to the usually high mutational burden in cancers [30-33]. However, we note that our model has been developed for normal tissue and does not account for chromosomal rearrangements in malignancies, which likely impact the estimation of mutation rates. It is an intriguing thought that early organ growth during development constitutes a very similar situation in which strand segregation is less effective within expanding stem cell populations and the increased rate of mutation accumulation early in life emerges as a natural consequence [34].

## Materials and Methods

We assume that homeostasis in a healthy adult human tissue is maintained by a constant pool of *N* stem cells. Each of these stem cells undergoes *n* cell divisions during a time interval Δ*t*. With each division, a stem cell non-randomly segregates DNA strands with a probability *p*. If *p* = 1 the ancestral strand will remain in the stem cell and the duplicated strand will be passed onto a daughter cell that becomes a non-stem cell, whereas *p* = 0.5 implies random strand segregation (i.e. no strand segregation), see Figure 1. We assume the probability *p* to be the same for all stem cells and don’t account for possible variation by for example specific mutations that would change strand segregation probabilities for individual stem cells. The non-ancestral duplicated strand inherits on average *µL* novel mutations, where *µ* is the mutation rate per base pair per cell division and *L* the length of the copied genome (e.g. *L* ≈ 6×10^9^ base pairs in humans). Throughout the manuscript we assume a constant mutation rate *µ*. In principal the mutation rate could depend on time explicitly, e.g. *µ* → *µ*(*t*). However, this would lead to non-linear dependencies, which is not supported by the currently available data, e.g. Figure 3 & 4. Thus assuming a constant mutation rate is retrospectively justified by the actual change of the mutational burden in human tissues.

It follows that for *n* cell divisions, the probability to segregate parental DNA strands *k* times is binomially distributed (*k* successes in *n* draws given a success probability of *p*)

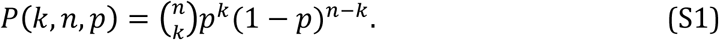

This implies that on average *E*[*k, n, p*] = *np* cell divisions do not induce additional mutations in stem cells. However, *n*(1 − *p*) cell divisions will increase mutational burden within a single stem cell lineage, each division by a random number *χ*, given by a Poisson distribution:

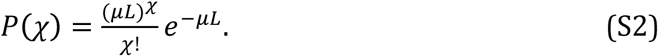

The mutational burden 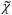 within a single stem cell lineage consequently increases by

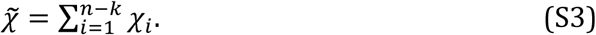

Exact expressions for the mutational burden 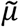 and variance σ^2^ for such distributions are known[35]. The mutational burden 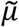 after *n* stem cell divisions is given by

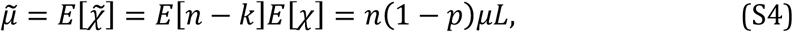

and the variance of the mutational burden σ^2^ is given by

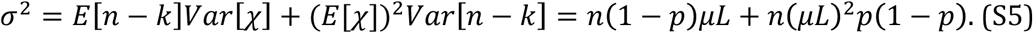

These expressions allow quantifying the change of the mutational burden as well as the change of the variance of the mutational burden after a number of Δ*n* divisions per stem cell

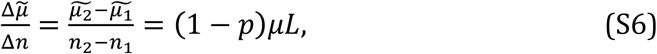

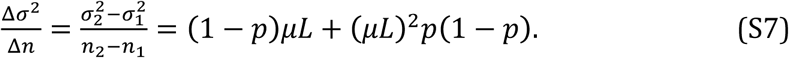

However, in actual data the number of stem cell divisions is unknown and change would be measured in time *t*. Assuming a constant rate of stem cell proliferations *λ* we can write Δ*n* = *λ* Δ*t*. This allows us to rewrite above equations for the change of the mean and the variance of the mutational burden over real time *t* via

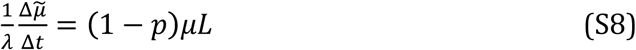

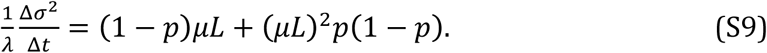

Importantly, both the change of the mutational burden 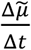 as well as the change of the variance of the mutational burden 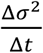 can be measured from human somatic mutation data, see Figure 3 & 4. Furthermore, equations (S8) and (S9) imply that the mutational burden as well as the variance of the mutational burden are expected to increase linearly with age in adult tissues. Even if strand segregation is highly effective, mutations still accumulate linearly with age. However, the rate of mutation accumulation is decreased by a factor of (1 − *p*). As we neither know the somatic mutation rate *µ* nor the stem cell proliferation rate *λ* with certainty, a linear increase in mutational burden with age at most suggests imperfect strand segregation (e.g. 0 ≤ *p* < 1). Importantly, the linear increase of both the mean and the variance in time is a result of the sum of Poisson distributed random variables and does by itself not imply the presence or absence of non-random strand segregation.

However, the ratio of variance and mean is independent of time *t* and the stem cell proliferation rate *λ*

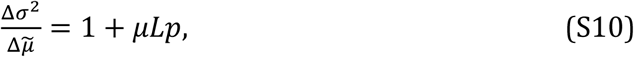

and therefore provides natural bounds for possible mutation rates per cell division *µ* and strand segregation probabilities *p* in human tissues, see SI Figure 1. Furthermore, rearranging equation (S8) and substituting 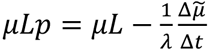 into equation (S10), the strand segregation probability *p* and the mutation rate *µ* disentangle, allowing us independent estimates via

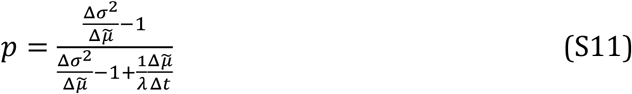

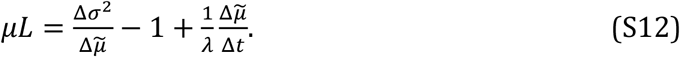

The relative change of the mutational burden and the variance variance of the mutational burden allow estimates of the mutation rate *µ* (per cell division) and the non-random strand segregation probability *p*. Estimating the mutation rate as well as the strand segregation probability, we need to measure the change of the mutational burden as well as the change of the variance of the mutational burden. This requires multiple measurements of the mutational burden within single cells of a single individual that ideally would be followed over time. This is unpractical and such data currently does not exist. We therefore measure the mutational burden and variance in multiple cells of multiple individuals of different ages. To calculate the variance and the mean of the mutational burden, we require at least 3 samples per individual, see Figure 3 & 4. For completeness we also show expressions for the mutation rate *µ* and p in dependence of stem cell proliferations *n*. They are given by

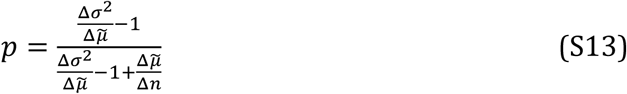

and

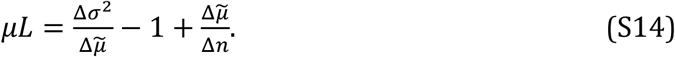

We recognize that our model is based on some assumptions and approximations. For example, telomeres, the protective ends of chromosomes, shorten with each cell division. Upon reaching a critically short telomere length, cells enter senescent. Senescence is not modelled in our model, however we argue that since this is likely to occur at very old ages [21,36], this process is unlikely to influence our results significantly.

## Acknowledgments

B.W. is supported by the Geoffrey W. Lewis Post-Doctoral Training fellowship. A.S. is supported by The Chris Rokos Fellowship in Evolution and Cancer and by Cancer Research UK (A22909). A.S. is also supported by the Wellcome Trust (202778/B/16/Z). This work was also supported by Wellcome Trust funding to the Centre for Evolution and Cancer (105104/Z/14/Z).

---

**SI Figure S1:**
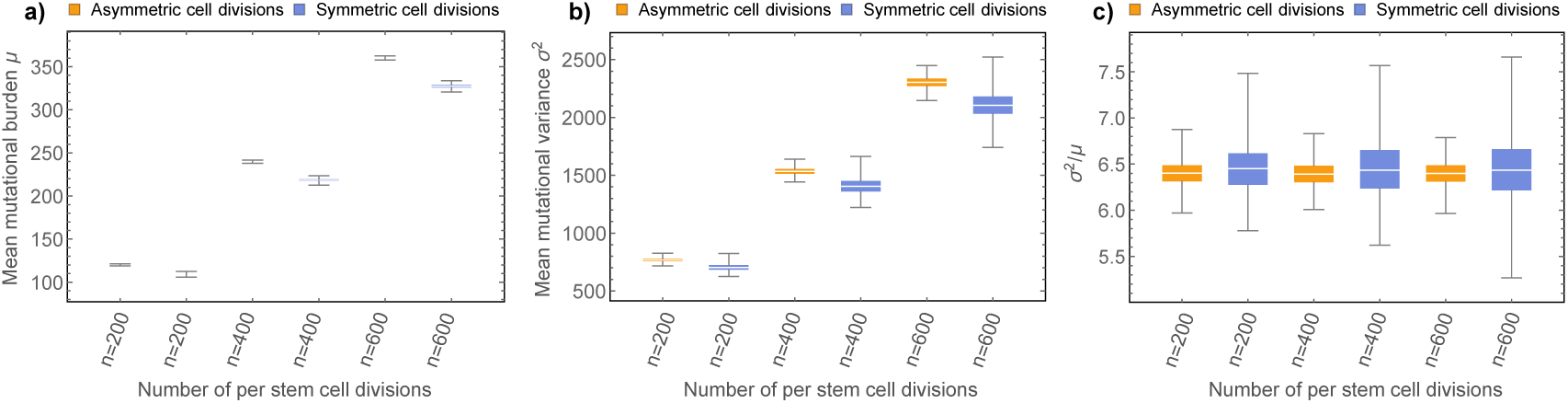
Strand segregation in terms of symmetric stem cell divisions. **a)** Mean mutational burden *µ*, **b)** mutational variance *σ*^2^, and **c)** the ratio of mutational variance and mutational burden *σ*^2^/*µ* for purely asymmetrically or a mix of symmetrically and asymmetrically dividing stem cells. Here we compare stochastic simulations for *N* = 5000 purely asymmetrically dividing stem cells with a strand segregation probability of *p* = 0.9 and stem cells with perfect strand segregation *p* = 1 but a fraction of 10% of stem cell divisions being symmetric differentiations followed by symmetric self-renewals. Both scenarios lead to a linear increase of mean and variance of mutational burden with minimal rate differences. However, as predicted, the ratio of variance and mean become time independent and are the same on average for both processes. However, the variance of the distribution of the ratio of the variance and mean increases with time for symmetric stem cell divisions but is approximately constant for asymmetric stem cell divisions. This effect might provide a future method to distinguish and quantitate the amount of symmetric self-renewal in human stem cell populations.

**SI Figure S2:**
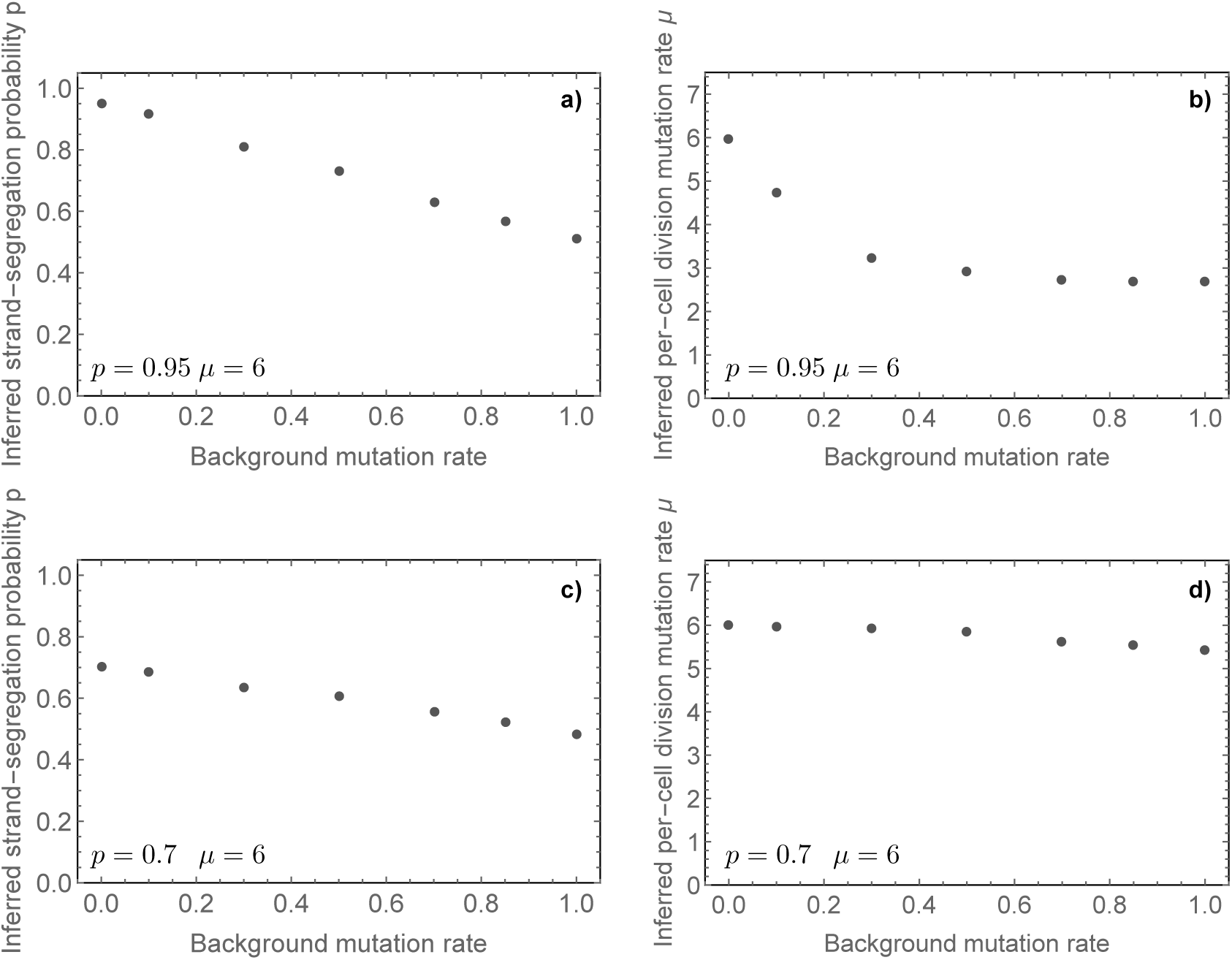
Influence of cell division independent background mutation rate on inference of non-random strand segregation probability and per-cell mutation rate. Plots **a)** to **d)** show the non-random strand segregation probability *p* and the per cell division mutation rate *µ* based on equations (S11) and (S12) inferred from stochastic simulations if we in addition allow for a constant cell-division independent mutation rate that influences both the ancestral and the duplicated DNA strand equally. In the upper panels **a)** and **b)** the underlying true parameters per cell division are *µ* = 6 and *p* = 0.95, whereas in the lower panels **c)** and **d)** we have *µ* = 6 and *p* = 0.7. If the background mutation rate is 0, we recover the original parameters. Both the non-random strand segregation probability *p* as well as the per cell division mutation rate *µ* are slightly underestimated for an increasing background mutation rate. Importantly, the non-random strand segregation probability is always underestimated and inferences become biologically meaningless (e.g. *p* < 0.5) for large background mutation rates. The actual data suggests high non-random strand segregation probabilities (see main text) and therefore implies small background mutation rates compared to cell division induced mutations.

